# FitHiChIP: Identification of significant chromatin contacts from HiChIP data

**DOI:** 10.1101/412833

**Authors:** Sourya Bhattacharyya, Vivek Chandra, Pandurangan Vijayanand, Ferhat Ay

## Abstract

Here we describe *FitHiChIP* (github.com/ay-lab/FitHiChIP), a computational method for identifying chromatin contacts among regulatory regions such as enhancers and promoters from HiChIP/PLAC-seq data. FitHiChIP jointly models the non-uniform coverage and genomic distance scaling of HiChIP data, captures previously validated enhancer interactions for several genes including *MYC* and *TP53*, and recovers contacts genome-wide that are supported by ChIA-PET, promoter capture Hi-C and Hi-C data. FitHiChIP also provides a framework for differential contact analysis as showcased in a comparison of HiChIP data we have generated for two distinct immune cell types.

---

Even though the invent of high throughput conformation capture techniques (e.g., Hi-C [1, 2], ChIA-PET [3]) has revolutionized the 3D genomics field, it has been challenging and extremely costly to generate kilobase resolution contact maps that allow for *de novo* identification of interactions among discrete regulatory elements [2]. Two new techniques that combine Hi-C with ChIP-seq, namely HiChIP [4] and PLAC-seq [5], show significant improvement over ChIA-PET [3] in direct profiling of regulatory (e.g., H3K27ac) and structural (e.g., cohesin) interactions with moderate sequencing depth (~200M reads) and in primary cells. However, at present, computational identification of functionally important subset of interactions/loops/contacts from this data remains difficult since these assays have specific biases and by design do not provide uniform coverage across the genome. Some of these difficulties have also hindered their use in detecting “differential interactions” that are due to specific changes in 3D looping (i.e., Hi-C component) and not due to changes in 1D coverage (i.e., ChIP-seq component). The original articles describing both assays [4, 5] (we use HiChIP to refer to both hereafter) use Hi-C-specific computational methods for analysis, which are not optimal for HiChIP data and hence suffer from low sensitivity (HiCCUPS [2]) or low specificity (Fit-Hi-C [6]). A recent tool *hichipper* provides a useful correction for 1D peak calling, but it suffers from low specificity in loop calling [7].

We have developed a versatile method, FitHiChIP (github.com/ay-lab/FitHiChIP), that identifies significant contacts from HiChIP data by: **i)** characterizing expected counts using a regression model that accounts for assay-specific biases as well as the scaling of contact probability with genomic distance **(see Methods)**, **ii)** computing statistical significance using a global background of expected counts, and **iii)** merging nearby loop calls using a connected component modeling to account for local density of interactions in order to improve specificity **(Fig. 1A Methods)**. FitHiChIP allows the users to either infer peaks from their HiChIP data or input a predefined reference set of ChIP-seq peaks **(Fig. 1A)**. The peak definitions are used to compute coverage bias values for each genomic window and to label pairs of windows according to their peak status, which are jointly used to infer either a stringent (S: peak-to-peak pairs only) or a loose background (L: peak-to-all pairs) of contact probability for each genomic distance. For the foreground, FitHiChIP also allows considering only peak-to-peak pairs, similar to most ChIA-PET pipelines, or including peak-to-nonpeak interactions for statistical assessment **(Fig. 1A)**. The resulting set of significant FitHiChIP loops provide for an intuitive way to perform differential analysis between cell types, which can also be coupled with HiChIP or ChIP-seq 1D coverage values to distinguish differential 3D looping from changes in loop visibility.

**Figure 1.**
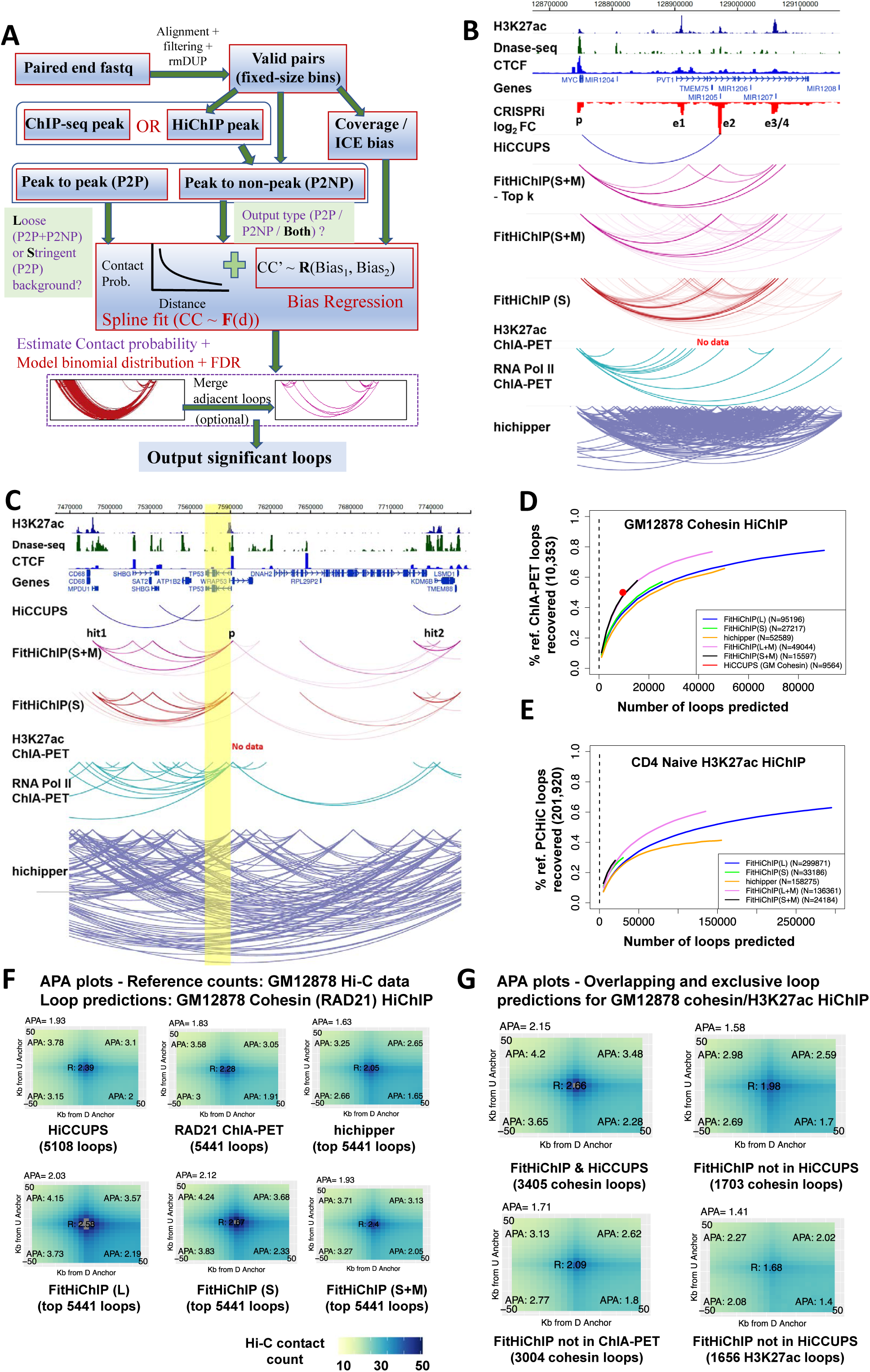
Identification of significant contacts by FitHiChIP. **(A)**: Overview of FitHiChIP pipeline. **(B)**: FitHiChIP accurately detects (H3K27ac HiChIP data from K562 cells) significant contacts between four distal distal enhancers (e1 to e4) and *MYC* promoter which are validated by CRISPRi [8]. Two of these loops are missed by HiCCUPS and and hichipper results are highly non-specific. **(C)**: For K562 H3K27ac HiChIP data, FitHiChIP reports interactions from the *TP53* promoter to two hit regions, located at ~100kb downstream and ~150kb upstream, respectively. Both of these loops are missed by HiCCUPS and and hichipper results are highly non-specific. **(D)**: FitHiChIP recovers higher percentage of reference ChIA-PET loops [12] compared to hichipper, for GM12878 cohesin HiChIP data [4]. **(E)**: FitHiChIP recovers higher percentage of reference promoter capture Hi-C loops on CD4 Naive T cells [14] compared to hichipper, when executed on CD4 Naive H3K27ac HiChIP data [11]. **(F)**: FitHiChIP loops are consistent with in-situ Hi-C data, as shown by higher APA scores of top-K significant loops (K=5441 for GM12878 cohesin HiChIP data) compared to hichipper. **(G)**: FitHiChIP loops among the top-K that are missed either by HiCCUPS calls on the same data (GM12878 cohesin or H3K27ac HiChIP [4, 11]) or by ChIA-PET (GM12878 cohesin [12]) exhibit considerably high APA scores suggesting they are supported by *in situ* Hi-C data. All of these results are for the distance range of 20kb-2Mb, includes peak to peak and peak to non-peak loops and use 5kb-binned HiChIP data.

To assess whether FitHiChIP connects distal enhancers to their experimentally validated target promoters from H3K27ac HiChIP data, we have compiled a list of loci for which functional data (e.g., CRISPRi) is available together with HiChIP data for the same cell line [8, 9, 10]. One gene with arguably the most comprehensive data is *MYC*, for which more than a megabase of surrounding region is tiled with sgRNAs for CRISPRi screen of cellular proliferation in K562 cells. This lead to identification of seven distal regions (e1–e7) ranging from 160kb to ~2Mb in distance to *MYC* [8]. All of these regions are identified as interacting with *MYC* promoter through RNA Pol II ChIA-PET [10] and Hi-C, hence are likely to inhibit proliferation through regulating *MYC* expression. For the ~400kb region with e1–e4, FitHiChIP identifies all four enhancer regions as interacting with *MYC* promoter from H3K27ac HiChIP data from K562 cells, whereas HiCCUPS (applied to H3K27ac HiCHIP) [11] only reports the interaction with e2 and hichipper non-specifically reports hundreds of interactions in this locus **(Fig. 1B)**. When sorted by significance, five out of top six FitHiChIP interactions either connect the *MYC* promoter to e1, e2 or e3-e4 (within the same bin) or connect enhancers to each other (e1/e2 to e3-e4). All methods capture the ~2Mb distal interactions between *MYC* and e5 or e6-e7. Similar CRISPRi data from the same work identifies no distal (>20kb) enhancer regions for *GATA1*. For another important gene, *TP53*, super-enhancer and broad domain analysis coupled with RNA Pol II ChIA-PET data in K562 cells identify two hit regions: one ~100kb downstream and one ~150kb upstream of the *TP53* promoter [10]. Using EpiSwitch baits, both regions are shown to interact with the *TP53* promoter, in addition to K562, in all CML patients (9/9) and nearly all normal subjects. FitHiChIP identifies both interactions with several other contacts in the domain, whereas HiCCUPS misses both and hichipper again reports hundreds of interactions **(Fig. 1C)**. Another two genes for which several CRISPRi and CRISPR KO experiments are carried out in K562 cells are *MYO1D* and *SMYD3* [9]. For both genes, FitHiChIP reports the interaction between the promoter and a *hub* enhancer with the highest impact on gene expression. Similar to above examples, HiCCUPS is too stringent to capture these interactions and hichipper is non-specific **(Supp. Figs. 1 2)**. Interestingly, both H3K27ac and RNA Pol II ChIA-PET [12, 10] also miss these two interactions.

**Figure 2.**
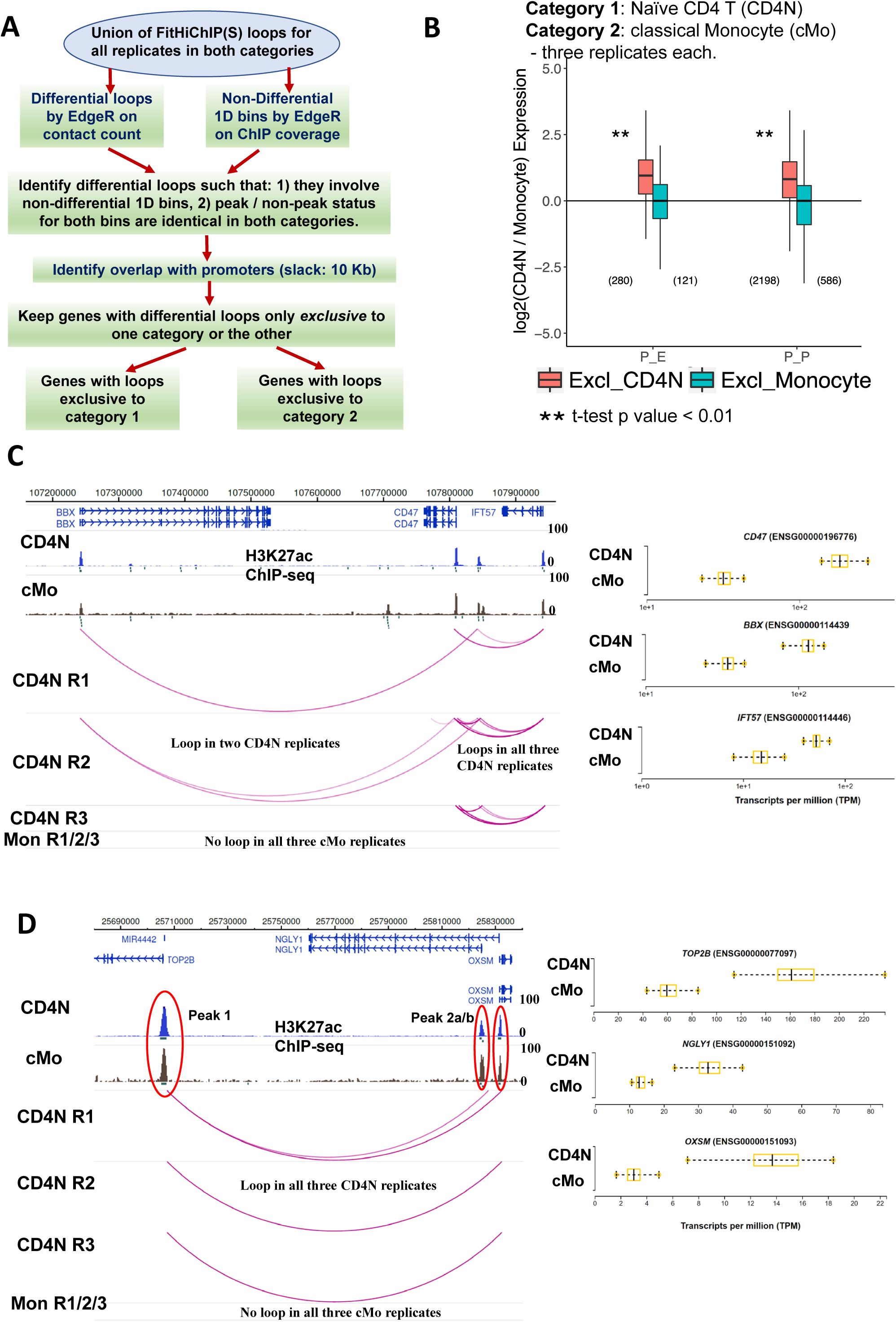
Identification of “3D differential” loops between two cell types using FitHiChIP. **(A)**: Schematic describing the identification of differential loops and further filtering of them to extract loops that are different due to 3D changes rather than 1D. **B**: 3D differential loops from promoters that are exclusive to one cell type generally relate to higher gene expression in that cell types, as observed by comparing three replicates of H3K27ac HiChIP data we have generated from Naive CD4 T cell (CD4N) and classical Monocyte (cMo). **C**: An example of 3D differential loops among the genes *IFT57*, *CD47* and *BBX*. The loop from *IFT57* to an intergenic enhancer and to *CD47* exist in all three replicates of CD4N, but in none of the cMo replicates. Another two loops from *CD47* to an enhancer and to *BBX* is present in two out of three CD4N replicates, but in none of the cMo replicates. Variability of expression for these genes in CD4N and cMo samples from 91 donors are shown as box plots as obtained from the DICE database [21]). **D**: Another example locus with a 3D differential loop involving the genes *TOP2B*, *NGLY1*, and *OXSM*. This loop is present in all three replicates of CD4N, but in none of the cMo replicates. Gene expression values for these genes in CD4N and cMo show all are expressed at higher levels in CD4N.

To systematically compare our method with existing tools and to evaluate the impact of different parameter selections, background models and normalization options in FitHiChIP performance, in the lack of large-scale and orthogonal gold standard data, we quantify the extent of concordance between statistical significance estimates from HiChIP data and other published cell type-matched contact data **(Supp. Tables 1 - 4)**. For each competing method or setting, we ask what fraction of reference loops reported by ChIA-PET (H3K27ac [12] and Pol2 [10] for K562 cells, RAD21 for GM12878 [12] and mouse embryonic stem cells [13]) or promoter capture Hi-C (naive CD4 cells [14]) are captured at differing number of loop calls from HiChIP data. In all cases FitHiChIP outperforms hichipper in both sensitivity and specificity **(Fig. 1D-E, Supp. Fig. 3)** with merging (M) of nearby interactions via 8-connectivity based connected component analysis [15] improving FitHiChIP’s specificity **(see Supp. Figs. 4, 5, and Methods** for details and parameters related to merging). These comparative results are robust to the choice of initial set of 1D peaks used **(Supp. Fig. 6)** and to the use of more stringent sets of reference loops **(Supp. Figs. 7, 8)**. In general, using ChIP-seq peaks as input rather than inferring 1D peaks from HiChIP data performs better in recovery of ChIA-PET or PCHiC loops **(Supp. Fig. 9)**. For inference of 1D peaks from HiChIP data, FitHiChIP and hichipper perform very similar in recovering a reference set of ChIP-seq peak calls **(Supp. Figs. 10, 11)** with hichipper background correction for restriction cut sites showing improvement for one case **(Supp. Figs. 10(a), 11(a))**. Finally, the regression-based correction for coverage bias differences by FitHiChIP results in better recovery of reference loops compared to matrix-balancing-based (e.g., ICE [16] which also introduces artifacts for several low coverage regions of HiChIP data) bias correction and to using no normalization **(Supp. Figs. 12, 13)**.

As exemplified by the above cases, in general, compared to FitHiChIP, HiCCUPS is too stringent and misses some of experimentally validated or ChIA-PET identified interactions whereas hichipper is highly non-specific and sometimes not informative. To further assess the significance estimations from each method by controlling for their differences in stringency, we turn to aggregated peak analysis (APA) of top-*k* predictions by utilizing high-resolution Hi-C data available for GM12878 cells [2]. HiCCUPS on GM12878 Cohesin HiChIP [4] as well as matching RAD21 ChIA-PET data [12] both report nearly 5000 loops within the suggested distance range of 150kb to 1Mb for APA analysis [17]. We sample similar number of top-*k* loops from FitHiChIP and hichipper and compare APA scores for each set. HiCCUPS and FitHiChIP loops show higher enrichment of center pixels as well as averaged APA score compared to both ChIA-PET and hichipper with hichipper performing worse **(Fig. 1F)**. Similar results hold when we repeat the same analysis with H3K27ac HiChIP data **(Supp. Fig. 14)**. Notably, cohesin HiChIP loops are stronger in enrichment compared to H3K27ac loops regardless of the loop prediction method. We then ask whether the interactions exclusively reported in top-*k* loops by FitHiChIP and not by either of the more stringent sets (HiCCUPS or ChIA-PET) still show enrichment in APA plots (i.e., supported by Hi-C data). For cohesin, nearly 3000 and 1700 loops exclusive to FitHiChIP when compared to HiCCUPS and ChIA-PET, respectively, both show strong APA scores and enrichment of center pixel, suggesting these loops are visible in the Hi-C data **(Fig. 1G Supp. Tables 6 - 7)**. Another 1656 loops exclusive to FitHiChIP on H3K27ac HiChIP data compared to HiCCUPS show a similar enrichment but to a lesser extent, consistent with overall lower APA scores compared to cohesin loops **(Fig. 1G Supp. Table 8)**. We also test FitHiChIP, HiCCUPS and hichipper results on GM12878 cohesin HiChIP data for four loci identified from Hi-C and validated by FISH to have strong CTCF-dependent long-range interactions [2]. All three methods capture the validated interactions from GM12878 data and almost none (except from one false positive for hichipper) identifies any interactions for the distance matched negative controls for FISH **(Supp. Figs. 15 - 18)**. Overall, these results suggest, beyond the set of strongest interactions that are captured independent of the loop calling method, there exists a considerable number of functionally validated and/or Hi-C/ChIA-PET/PCHiC supported HiChIP loops, which are identified exclusively by FitHiChIP.

Next, we use FitHiChIP loop calls as basis for differential chromatin contact analysis for HiChIP data with replicates. First, we have performed H3K27ac ChIP-seq and have profiled H3K27ac-mediated contacts in naive CD4 T cells (denoted as CD4N) and classical monocytes (denoted as cMo) sorted from leukapheresis samples of three blood donors (replicates) using the HiChIP protocol of [11] with modifications **(see Methods and Suppl. Table 5)**. To avoid multiple testing on the whole contact map, we perform differential analysis on a set of contacts that are deemed statistically significant in at least one replicate of one cell type **(Fig. 2A Supp. Fig. 19)**. Out of ~56k tested, nearly ~ 21k loops are called as differential by edgeR [18] (adjusted pval < 0.05 and fold change >2) and ~8k of these overlap with a promoter on at least one end. A major portion of these “differential loops” are between regions with altering H3K27ac levels (either from ChIP-seq or HiChIP coverage) across the two cell types and hence are likely changes in the visibility of loops by HiChIP rather than changes in the looping in 3D. To distinguish these two modes, we perform another differential analysis on the corresponding ChIP-seq coverage values at 5kb resolution using edgeR. We extract the set of differential loops (CD4N vs cMo) that do not overlap with a differential ChIP-seq bin (adjusted pval < 0.05 and fold change >2) on either side and also keep their peak/non-peak status unchanged both sides **(see Methods)**. This provides us with 2036 promoter loops which are not substantially different in 1D but are significantly different in 3D (i.e., 3D differential). We further extract promoters that only have CD4N or cMo exclusive loops, and thus obtain 2219 distinct genes (multiple promoters can overlap one loop) with a differential loop to another promoter (P-P) or an enhancer element (P-E). For both categories, the expression of genes are, in general, higher for the cell type with the loop (statistically significant for CD4N), suggesting 3D differential loops are relevant for gene expression changes across cell types **(Fig. 2B Supp. Fig. 19)**. An example for this phenomenon is the *CD47* which has nearly identical ChIP-seq tracks for CD4N and cMo (aside from several non-interacting cMo exclusive peaks) but only has significant and 3D differential P-P and P-E loops in CD4N **(Fig. 2C)**. Interestingly, all three genes (*BBX, CD47, IFT57*) in this ~700k locus show at least 3 fold higher gene expression in CD4N compared to cMo (http://dice-database.org). Both *CD47* and *IFT57* are known regulators of T cell activation and recently it has been shown that the two have correlated expression in T cells [19]. *CD47* is also known as the “don’t eat me” signal to macrophages in the context of cancer [20]. Another example is the ~200k *TOP2B* locus that has identical ChIP-seq coverage and peak calls across CD4N and cMo but has a P-P loop that connects all three genes in each replicate of CD4N and none of cMo **(Fig. 2D)**. *TOP2B*, a DNA topoisomerase is involved in relief of torsional stress during transcription and replication, as well as the two other genes all encode for important enzymes and are expressed at significantly higher levels in CD4N (P-P loop present) compared to cMo. Several other examples of 3D differential loops between these two cell types are shown in **Supp. Figs. 20 and 21**. These results suggest that even when controlled for 1D coverage differences, the differential loops identified from HiChIP data by our framework have a significant impact on gene expression for several genes. It is important to note that there also are cases where loop existence does not correlate with higher gene expression.

Overall, FitHiChIP provides an empirical null-based, flexible method for statistical significance estimation and differential loop calling from HiChIP data. Functional relevance of loops reported by our method is evident from the above discussed case-by-case and systematic analyses and from a genome-wide quantification that the genes with higher number of strong enhancer loops have higher expression **(Supp. Figs. 22 - 24)**. Our method also provides several options and modes for the user while keeping the number of tunable parameters to a minimum. An important feature of FitHiChIP is its ability to call loops between peak and non-peak regions. Such loops, which are generally discarded by previous methods including those for ChIA-PET, are at least equally supported by PCHiC loops as compared to peak to peak loops from H3K27ac HiChIP data **(Supp. Fig. 25)**. FitHiChIP is fast and scalable to 1 kb resolution even though all the analyses presented here used 5 kb bins. We believe FitHiChIP is a critical step in thoroughly exploring the rich data from HiChIP assay by facilitating its interpretation and analysis.

## Acknowledgements

We would like to thank Abhijit Chakraborty, Arya Kaul and all members of the Ay lab for their input and helpful comments on the tool development. We are also grateful to Benjamin Schmiedel, Gregory Seumois for their help in cell sorting, ChIP-seq and sequencing and all investigators of the R24 resource grant (R24AI108564) and the donors of leukapheresis samples. This work was funded in part by NIH/NIGMS award R35 GM128938 to F.A.

## Author contributions

S.B. and F.A. conceived the project and designed the method. S.B. implemented the software under supervision of F.A. V.C. carried out the HiChIP experiments. S.B and F.A. conducted the computational experiments, interpreted the results and drafted the manuscript with input from V.C. and P.V. All authors read and approved the final version of the manuscript.

## Methods

### Links to Epigenome Browser with all results visualized

- FitHiChIP GM12878 cohesin [4] session link: http://epigenomegateway.wustl.edu/browser/?genome=hg19&session=2r3JhBEXme&statusId=1665512412
- FitHiChIP GM12878 H3K27ac [11] session link: http://epigenomegateway.wustl.edu/browser/?genome=hg19&session=D5MTGCGTlo&statusId=1459152954
- FitHiChIP K562 H3K27ac [11] session link: http://epigenomegateway.wustl.edu/browser/?genome=hg19&session=SiANguifsa&statusId=2018280635
- FitHiChIP CD4 Naive H3K27ac [11] session link which also includes promoter capture Hi-C tracks [14]: http://epigenomegateway.wustl.edu/browser/?genome=hg19&session=m6RgEzdwqD&statusId=1181364500
- FitHiChIP mESC Cohesin [4] session link: http://epigenomegateway.wustl.edu/browser/?genome=mm9&session=qhIIdfNe4s&statusId=417062823
- FitHiChIP for our newly generated HiChIP and ChIP-seq data from primary naive CD4 T cells and classical monocytes (three replicates each) and differential analysis session link which also includes promoter capture Hi-C tracks [14]: http://epigenomegateway.wustl.edu/browser/?genome=hg19&session=HrTGl59KNa&statusId=1439871467

### Datasets used

We have used published HiChIP datasets **(Suppl. Table 1)** [4, 11] of four cell types: GM12878, K562 and Naive CD4 T cells (reference genome hg19); mouse embryonic stem cells (reference genome mm9), with two different proteins or histone marks of interest (cohesin as profiled either by RAD21 or Smc1/3 and H3K27ac). We have also generated HiChIP datasets on two immune cell types: Naive CD4 T cells and classical monocytes, using an antibody against H3K27ac for IP (**Suppl. Table 5**). Each of these datasets consist of three replicates (different donors). HiCCUPS loops (**Suppl. Table 2**) computed on the published HiChIP datasets [4, 11] are used for benchmarking FitHiChIP. ChIA-PET loops (**Suppl. Table 3**) are obtained from two previous publications [12, 13]. We have also used Promoter Capture Hi-C loops [14] on Naive CD4 T cells (using ChiCAGO [22] score ≥ 5 to call significant loops) as a reference (**Suppl. Table 4**). FitHiChIP loops reported in this study employ 5kb binning. ChIA-PET and hichipper loops are applied fixed size (=5 Kb) binning, by mapping them to the nearest 5 Kb bins. HiCCUPS loops contain a mixture of 5kb (mainly) and 10kb resolution.

### Datasets generated

HiChIP is performed as described in [11], with slight modifications. Briefly, 5× 10^6^ sorted cells are formaldehyde fixed, digested with MboI and biotinylated nucleotides are inserted using Klenow. After ligation, sonication is performed using Covaris and immunoprecipitation is performed using H3K27ac polyclonal antibody - Premium (Diagenode). Post ChIP, DNA marked by biotin are immobilized using MyOne Streptavidin C1 DynaBeads (Invitrogen) and is pulled down with streptavidin beads. Bead bound purified DNA is end repaired, adenine tailed, and ligated to adaptors. The immobilized HiChIP libraries are amplified using NEBNext multiplex oligoes for Illumina (New England Biolabs) with 10-12 PCR amplification cycles. Amplified library is size selected using AMPure XP beads to isolate DNA ranging from 300 to 750 bp in size and then sequenced.

### Preprocessing HiChIP data

Input paired fastq reads are processed through HiC-Pro pipeline [23] for their alignment using Bowtie2 [24], assignment to the restriction fragments (MboI), filtering by their orientation [25], and removal of duplicates using Picard tools [26]. Filtered reads are binned at fixed size (default 5kb) to generate the contact map/matrix. Individual bins are tested for their overlap with given set of peaks, and are termed either as a *peak bin* or a *non peak bin*, respectively.

### Types of interactions reported

FitHiChIP reports, by default, *peak-to-all* (both peak-to-peak and peak-to-nonpeak) interactions. It also supports reporting specifically peak-to peak, peak-to-nonpeak, or even all-to-all (interactions between every pair of bins simiar to Hi-C) loops. FitHiChIP considers interactions within a given distance range (default range 20 Kb - 2 Mb) which can be specified by the user.

### Inferring peaks from HiChIP data

For the purposes of calling 1D peaks from HiChIP data when ChIP-seq peaks are not avialable, we have tested different combinations of following four sets of reads generated from the HiC-pro pipeline: 1) dangling end (DE), 2) self-cycle (SC), 3) re-ligation (RE), and 4) CIS ‘short-range’ (<1 Kb) valid (V) reads (after duplicate removal). We have used MACS2 [27] on reads from each combination with the following parameters “-q 0.01–extsize 147 –nomodel” (suggested in hichipper [7]) to infer corresponding set of peaks. In addition, we have also used the peaks inferred from hichipper using DE + SC reads, with its custom background correction that models the decay of the number of reads according to the distance to the nearest restriction enzyme cut site.

### Spline fit for modeling the contact decay with increasing genomic distance

As suggested in FitHiC [6], FitHiChIP performs equal occupancy binning of input reads pairs into variable-sized genomic distance bins. If *N* = number of interacting locus pairs, *C* = total contact count, *M* = number of equal occupancy bins, *n*_*j*_ = number of locus pairs for individual bins *j* (1 ≤ *j* ≤ *M*), *S*_*j*_ = sum of contact counts for these *n*_*j*_ pairs in bin *j*, and *I*_*j*_ = average interaction distance for all *n*_*j*_ locus pairs within *j*, then average contact probability of bin *j* is *p*_*j*_ = (*S*_*j*_*/n*_*j*_)*/C*. The points (*p*_*j*_, *I*_*j*_) are used for spline fitting to model the distance decay. FitHiChIP can report either peak-to-all loops or peak-to-peak loops and can use either set of loops for learning the background and spline fitting regardless of the output mode. FitHiChIP(S) (S for stringent, also denoted as P2P=1) uses only peak-to-peak loops as background, whereas FitHiChIP(L) (L for loose, also denoted as P2P=0) employs background from peak-to-all loops. The stringent background reports much higher background probability of interaction and, hence, more conservative p-value/q-value estimates.

### Bias regression

FitHiChIP either uses coverage normalization (separately for peaks and non-peaks), or estimates bias using ICE [16]. For each individual equal occupancy bins, FitHiChIP uses each locus pair and its raw contact count, bias values, as well as the expected contact count estimated from the spline fit, to model a regression and to estimate the overall expected contact probability for that specific locus pair. If *C*_*I j*_ is the spline fitted contact count corresponding to the average interaction distance *I*_*j*_ for a bin *j*, *B*_1_ and *B*_2_ denote the bias vectors for the interacting bins of the set of locus pairs within the bin *j*, and *K* is the set of raw contact counts for these locus pairs, then the linear regression model is defined as log(*K*) = *R* (*log*(*B*_1_)*, log*(*B*_2_)*, log*(*C*_*I*_*j*)). Such regression produces the expected contact count 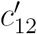for individual locus pairs (*l*_*1*_, *l*_*2*_). The expected contact probability *p* for such a locus pair becomes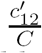. Binomial distribution on the contact probability generates a p-value which is then corrected by the Benjamini-Hochberg procedure [28] for multiple testing. The resulting q-value is reported as the statistical significance. An FDR threshold of 0.01 is used to report the set of significant interactions.

### Merging adjacent loops

Two loops are *adjacent* if their constituent bins are either adjacent or equal. If we represent each loop L between two 5 Kb bins x and y as a non-zero element in a matrix of bins M, finding a set of mutually adjacent loops becomes finding a non-trivial connected component [15] of non-zero elements within M, using the 8-connectivity rule. We use the Python package *networkx* [29] to find such components. For each component, either its most significant loop (lowest q-value) can be used (denoted as the *MIN* approach).However, this method discards a high number of loops for larger components. Therefore, we select a set of loops within the connected component which are mutually separated by at least B bins in both sides (thus separated by W = BxB window in the 2D matrix). This technique is termed as *iterative merging*. We have experimented with B = 2, 5 and 10, and found that B = 2 performs the best when evaluated by the recovery plots.

### Computing overlap with reference HiCCUPS or ChIA-PET loops

As some of the reference HiCCUPS loops employ 10 Kb binning, we have allowed 5 Kb slack for computing overlaps between HiCCUPS (or ChIA-PET) with FitHiChIP and hichipper loops. So, two interactions overlap if their respective bins lie within 5 Kb of each other. In each iteration, top X loops of FitHiChIP (or hichipper) and K number of reference HiCCUPS (or ChIA-PET) loops are tested for overlap. Here, X starts from K and is gradually incremented with a fixed step size. FitHiChIP loops are sorted by decreasing order of statistical significance (increasing q-value), while hichipper loops are sorted by decreasing order of the last column reported.

### Computing overlap with reference promoter capture Hi-C loops

For both FitHiChIP and hichipper, their promoter specific loops (loops whose at least one end overlaps with a reference TSS site with an allowed slack of 5 Kb) are used for computing overlap with the reference promoter capture Hi-C loops. Either the complete set of reference PCHi-C loops, or only the top 50K loops (in terms of higher significance, measured by ChICAGO [22] score) are used for comparison.

### Aggregate Peak Analysis (APA)

Reference Hi-C contact matrices (binned at 5 Kb) for different cell lines like GM12878, K562 [2] are first normalized by ICE [16]. We use such normalized Hi-C contact counts (mapped with respect to FitHiChIP or other reference loops) of 50 Kb up and downstream of individual interacting loci (as suggested in MANGO [17]) to compute the APA score. For 5 Kb binning, a contact matrix *M* of 21×21 dimension is thus generated, whose central entry corresponds to individual interacting locus pairs. Loops within a distance between 150 Kb and 1 Mb (as suggested in [17]) are only considered for APA analysis. The *APA score* [17] (displayed on top of each of the APA plots) is the ratio of the central pixel and the mean of pixels 15-30 Kb downstream of the upstream loci and pixels 15-30 Kb upstream of the downstream loci.

As the number of interactions reported by FitHiChIP or hichipper are substantially higher than HiCCUPS or ChIA-PET loops, we use top-K loops of FitHiChIP or hichipper (in terms of higher statistical significance) for APA analysis, where K is the number of loops reported by HiCCUPS or ChIA-PET.

Considering the APA score of overlapping and exclusive loops between FitHiChIP and the reference interactions, suppose K is the number of reference (HiCCUPS or ChIA-PET or hichipper) loops within the distance range 150 Kb - 1 Mb. Then the top-K loops of FitHiChIP (within the same distance range) are considered for overlap analysis. Overlapping and exclusive set of loops for either FitHiChIP or reference categories are separately analyzed for their APA scores. As mentioned previously, a slack of 5 Kb is allowed to compute the overlap.

### Gene expression versus the number of strong enhancers associated with promoters

We use the 15-state gene annotations provided in ChromHMM [30] for GM12878 and K562 cell lines, to determine the interactions between promoters and strong enhancers. Gene expression for GM12878 cell line is obtained from www.encodeproject.org (experiments/ENCSR000COR), while gene expression for K562 cell line is obtained from (experiments/ENCSR000CPS). Promoters are divided in five groups of 20 percentile each, based on the number of strong enhancers they interact as determined by the method to process HiChIP or by the ChIA-PET data.

### Differential analysis of FitHiChIP loops

We consider the union of significant (FDR < 0.01) loops generated by FitHiChIP(S) for the given pair of categories with *M* and *N* number of replicates. The package edgeR [18] is used on the integer ratio of raw contact count and the expected contact count (from the bias regression module) to get the initial set of differential loops. Only the P-P or P-E differential loops are processed further. These loops are differential either due to the difference in their 1D (ChIP) coverage or their 3D (HiChIP) coverage. Therefore, we use edgeR to derive the fixed size (here 5 Kb) bins which are non-differential in terms of their ChIP (1D) coverage (computed using the merged ChIP-seq alignment from all the replicates). Differential loops which involve only the non-differential bins are processed. As the HiChIP loops are of type either peak-to-peak or peak-to-all, one reason for the formation of a differential loop is the difference in peak status of a particular bin (i.e. it is peak in one category and non-peak in other). Thus, we have considered only those loops whose interacting bins have identical peak status in both categories. The resulting loops are differential solely due to the difference in HiChIP coverage. We further extract the differential loops which are exclusive (present in one or more replicates) in one category (and not present in the other category). If the given categories correspond to different cell types (such as Naive CD4 T cell and classical Monocyte), we check whether the exclusive loops (of one category) also exhibit statistically significant higher expression compared to the other category (significance is measured by sample t-test [31], with p-value threshold of either 0.05 or 0.01).

